## Supplementary material for "Intellectual disability-associated disruption of O-GlcNAcylation impairs neuronal development and cognitive function in *Drosophila*"

**Intellectual disability-associated disruption of O-GlcNAc cycling impairs cognition in *Drosophila***

**This PDF file includes:**

Figures S1 to S4

Tables S1 and S2

SI References

**Other supplementary materials for this manuscript include the following:**

Table S3

Table S4

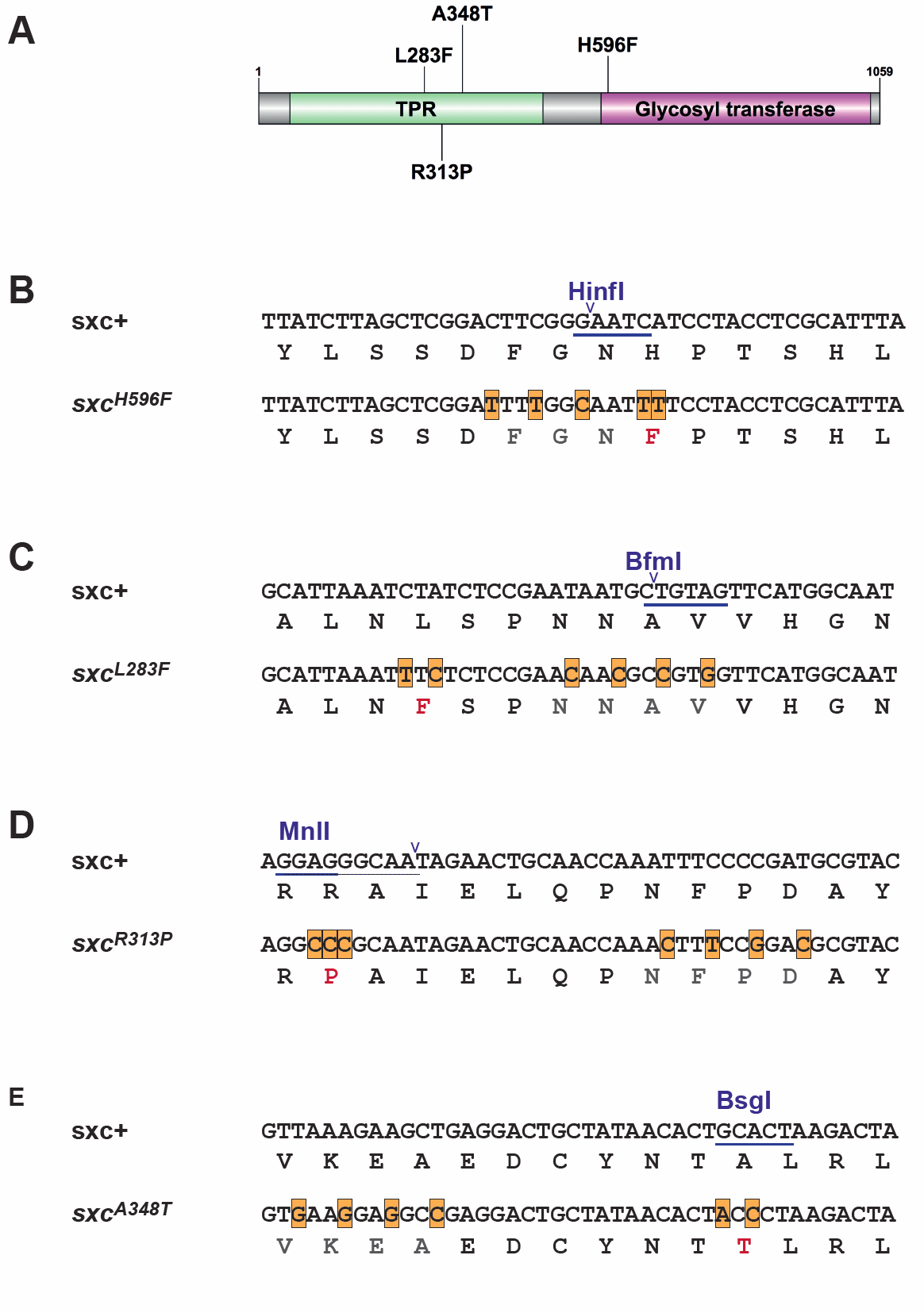
**Fig. S1.** **Figure S1. Generation and characterization of *sxc^H596F^*, *sxc^L283F^*, *sxc^R313P^* and *sxc^A348T^* alleles.**

(A) Schematic representation of Drosophila sxc protein showing the location of H596F, L283F, R313P and A348T mutations; purple tetratricopeptide repeat (TPR) domain, green glycosyl transferase (GT) domain.
(B) **– (E)** Sequences of genomic DNA of wild type, sxcH596F, sxcL283F, sxcR313P and sxcA348T Drosophila alleles. The missense mutation and additional silent mutations are highlighted. The restriction digestion sites used for genotyping are shown.

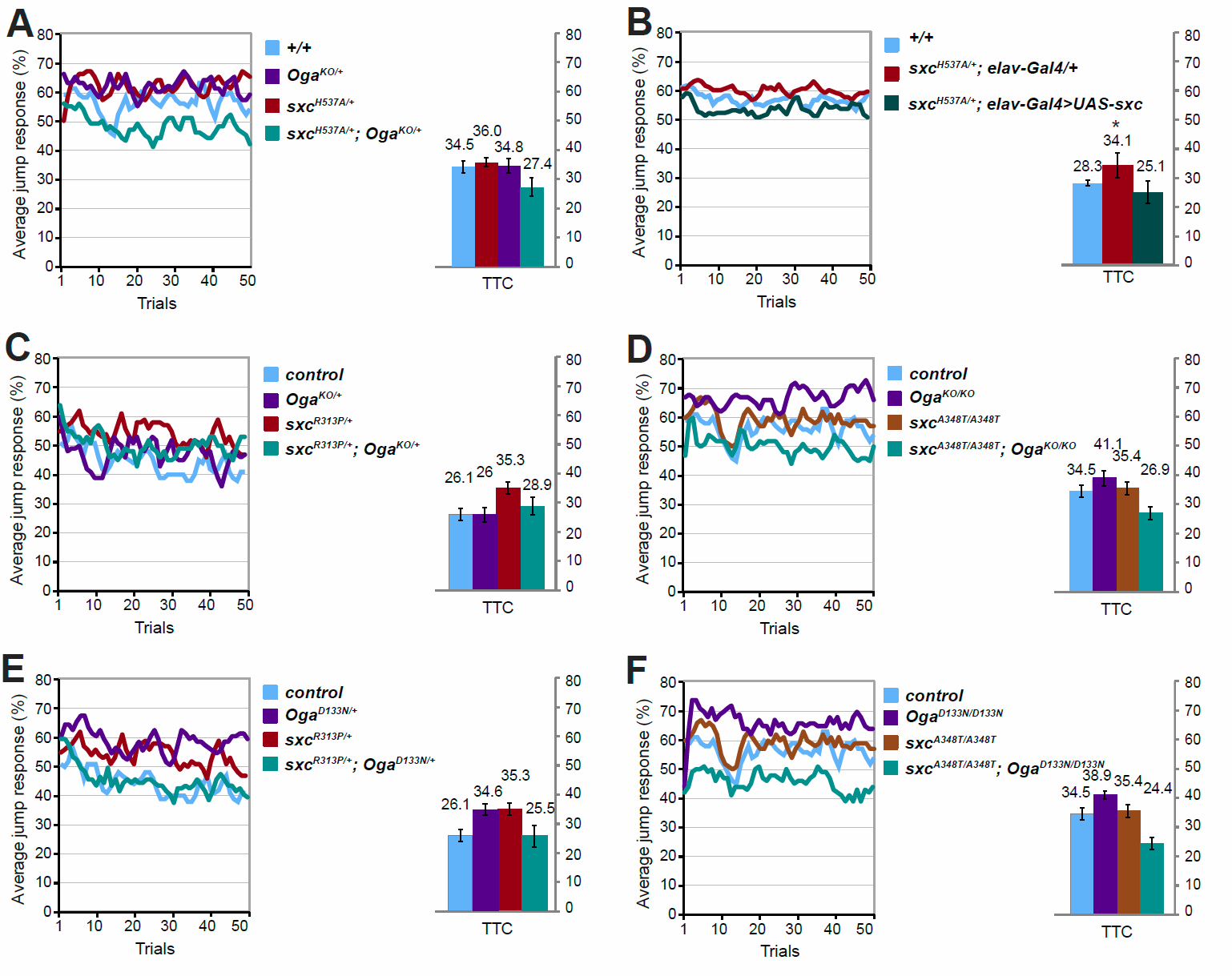

**Fig. S2.** **Jump responses in the fatigue assay**

In the fatigue assay, jump responses were induced with 50 light-off pulses with 5 s interval between pulses that prevents habituation. The jump response is presented as % of jumping flies in each light-off trial. The mean number of trials that flies needed to reach the no-jump criterion (Trials To Criterion, TTC) ± SEM is presented.

(A) Jump response of the *sxc^H537A/+^; Oga^KO/+^* flies (N = 85, mean TTC ± SD: 27.4 ± 8, in cyan) remains high throughout the entire course of the experiment, similar to control flies (*+/+*, N = 85, mean TTC ± SD: 34.5 ± 5, *p*_adj_ = 0.14, in blue) demonstrating that restored habituation in *sxc^H537A/+^; Oga^KO/+^* flies (**Figure 1B**) is not confounded by fatigue.
(B) Jump response of the *sxc^H537A/+^; elav-Gal4>UAS-sxc* flies (N = 52, mean TTC ± SD: 25.1 ± 7.9, in green) remains high throughout the entire course of the experiment, similar to control flies (*+/+*, N = 55, mean TTC ± SD: 28.3 ± 2.3, *p*_adj_ = 0.128, in blue) demonstrating that restored habituation in *sxc^H537A/+^; elav-Gal4>UAS-sxc* flies (**Figure 1D**) is not confounded by fatigue.
(C) Jump response of the *sxc^R313P/+^; Oga^KO/+^* flies (N =73, mean TTC ± SD: 28.9 ± 7.6, in cyan) remains high throughout the entire course of the experiment, similar to control flies (*+/+*, N = 84, mean TTC ± SD: 26.1 ± 5.1, *p*_adj_ = 1, in blue) demonstrating that restored habituation in *sxc^R313P/+^; Oga^KO/+^* flies (**Figure 4C**) is not confounded by fatigue.
(D) Jump response of the *sxc^A348T/A348T^; Oga^KO/KO^* flies (N = 78, mean TTC ± SD: 26.9 ± 5.1, in cyan) remains high throughout the entire course of the experiment, similar to control flies (*+/+*, N = 85, mean TTC ± SD: 34.5 ± 5, *p*_adj_ = 0.31, in blue) demonstrating that restored habituation in *sxc^A348T/A348T^; Oga^KO/KO^* flies (**Figure 4D**) is not confounded by fatigue.
(E) Jump response of the *sxc^R313P/+^; Oga^D133N/+^* flies (N = 83, mean TTC ± SD: 25.5 ± 9.2, in cyan) remains high throughout the entire course of the experiment, similar to control flies (*+/+*, N = 84, mean TTC ± SD: 26.1 ± 5.1, *p*_adj_ = 1, in blue) demonstrating that restored habituation in *sxc^R313P/+^; Oga^D133N /+^* flies (**Figure 4E**) is not confounded by fatigue.

Jump response of the *sxc^A348T/A348T^; Oga^D133N/D133N^* flies (N = 63, mean TTC ± SD: 24.4 ± 5.1, in cyan) remains high throughout the entire course of the experiment, similar to control flies (*+/+*, N = 85, mean TTC ± SD: 34.5 ± 5, *p*_adj_ = 0.15, in blue) demonstrating that restored habituation in *sxc^A348T/A348T^; Oga^D133N/D133N^* flies (**Figure 4F**) is not confounded by fatigue. * *p*_adj_<0.1, based on lm analysis with Bonferroni-Holm correction for multiple comparisons. Complete list of p-values and summary statistics is provided in **Table S3**.

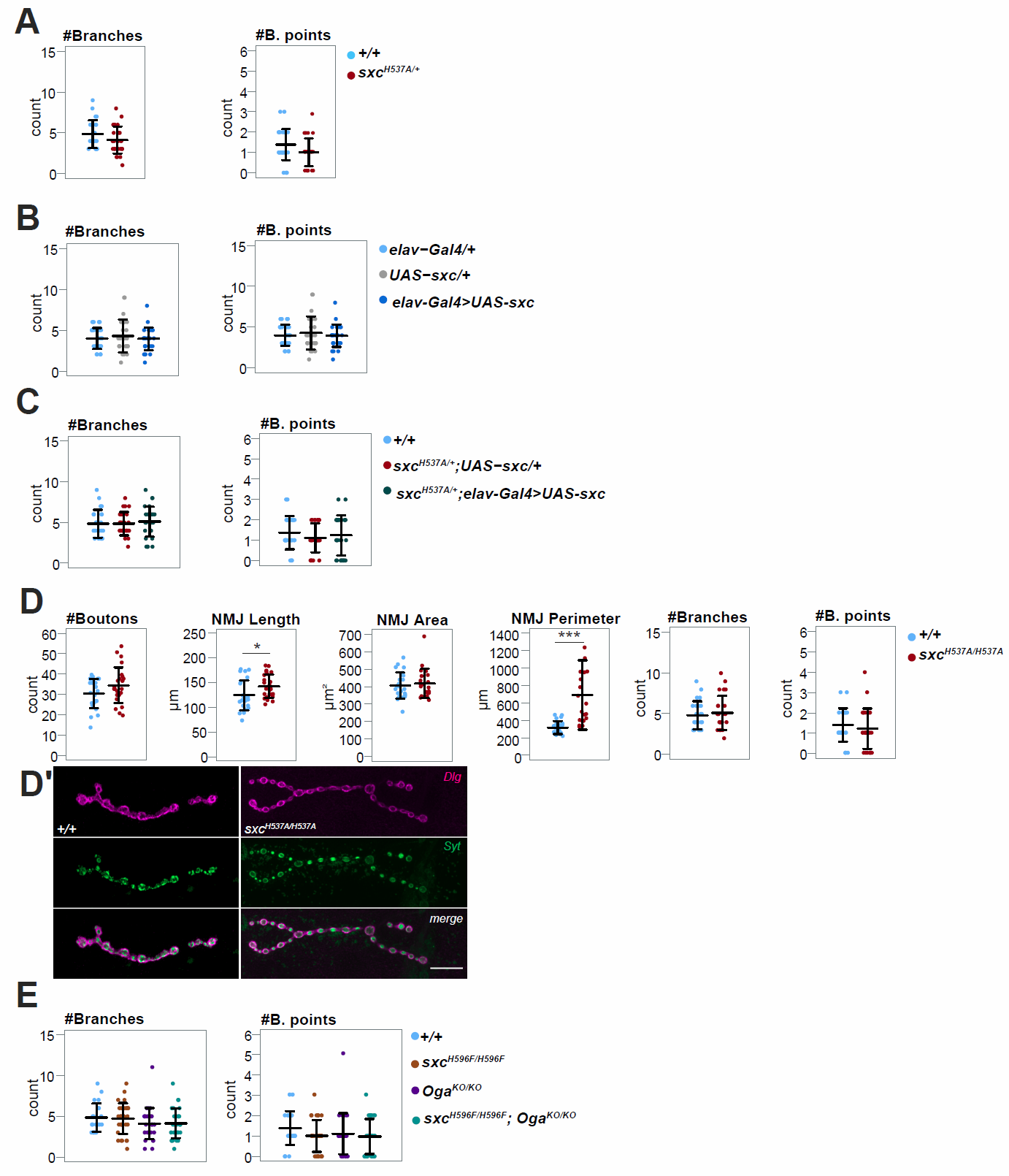

**Fig. S3.** **NMJ of *sxc^H537A/H537A^* mutant and NMJ branching morphology**

(A) Number of synaptic branches and branching points in *sxc^H537A/+^* larvae (N = 29, in red) is not significantly different from the genetic background control larvae (*+/+*, N = 24, branches: *p* = 0.1439, branching points: *p* = 0.05648, in blue). P-values are based non-parametric Wilcoxon test analysis.
(B) Number of branches and branching points is not affected in *elav-Gal4>UAS-sxc* larvae (N = 29, in dark blue) compared the *elav-Gal4/+* larvae (N = 30, branches: *p*_adj_ = 0.8702, branching points: *p*_adj_ = 0.8488, in light blue) and to *UAS-sxc/+* larvae (N = 26, branches: *p*_adj_ = 0.8294, branching points: *p*_adj_ = 0.2689, in grey).
(C) Branches and branching points are not affected in *sxc^H537A/+^; UAS-sxc/+* larvae (N = 28, in red) compared to the control larvae (*+/+*, N = 28, branches: *p*_adj_ = 0.7121, branching points: *p*_adj_ = 0.2979, in blue). *sxc^H537A/+^; elav-Gal4>UAS-sxc* larvae (N = 29, in green) do not show any changes in number of branches and branching points compared to the *sxc^H537A/+^; UAS-sxc/+* larvae (branches: *p*_adj_ = 0.4097, branching points: *p*_adj_ = 0.5928) and control larvae (*+/+*; branches: *p*_adj_ = 0.4301, branching points: *p*_adj_ = 0.6927).
(D) *sxc^H537A/H537A^* larvae have significantly increased NMJ length (N = 25, *p* = ) and perimeter (N = 21, *p* = , in red) compared to their genetic background control (*+/+*, N = 26, in blue) but not significantly different number of boutons (*p* = 0.085), NMJ area (*p* = 0.618), number of branches (*p* = 0.691) and branching points (*p* = 0.371). * *p*<0.05, *** *p*<0.001. P-values for boutons, length, area and perimeter are based on one-way ANOVA. P-values for branches and branching points are based on non-parametric Wilcoxon test analysis.

Branches and branching points are not affected in *sxc^H956F^* larvae (N = 31, branches: *p*_adj_ = 1, branching points: *p*_adj_ = 0.5, in brown), *Oga^KO^* larvae (N = 30, branches: *p*_adj_ = 0.75, branching points: *p*_adj_ = 0.51, in purple), and *sxc^H596F^; Oga^KO^* larvae (N = 30, branches: *p*_adj_ = 0.75, branching points: *p*_adj_ = 0.5, in cyan) compared to the genetic background control larvae (*+/+*, N = 28, in blue). *sxc^H596F^; Oga^KO^* larvae do not show a significant change in number of branches and branching points compared to the *sxc^H956F^* larvae (branches: *p*_adj_ = 0.75, branching points: *p*_adj_ = 1). Data presented as individual data points with mean ± SD. P-values are based on Kruskal-Wallis test with Wilcoxon pairwise test for multiple comparisons. Complete list of p-values and summary statistics is provided in Table S3.

(D’)	Representative NMJs of genetic background control (*+/+*) and *sxc^H537A/sxcH537A^* wandering	third instar larvae labeled with anti-discs large 1 (*Dlg*, magenta) and anti-synaptotagmin	(*Syt*, green). *Scale bar*, 20µm. The quantitative parameter values of the representative images (*+/+ | sxc^H537A/sxcH537A^*): #Boutons (26 | 30), Length (96.2 | 169.8), Area (415.9 | 464.3), Perimeter (258.4 | 445).

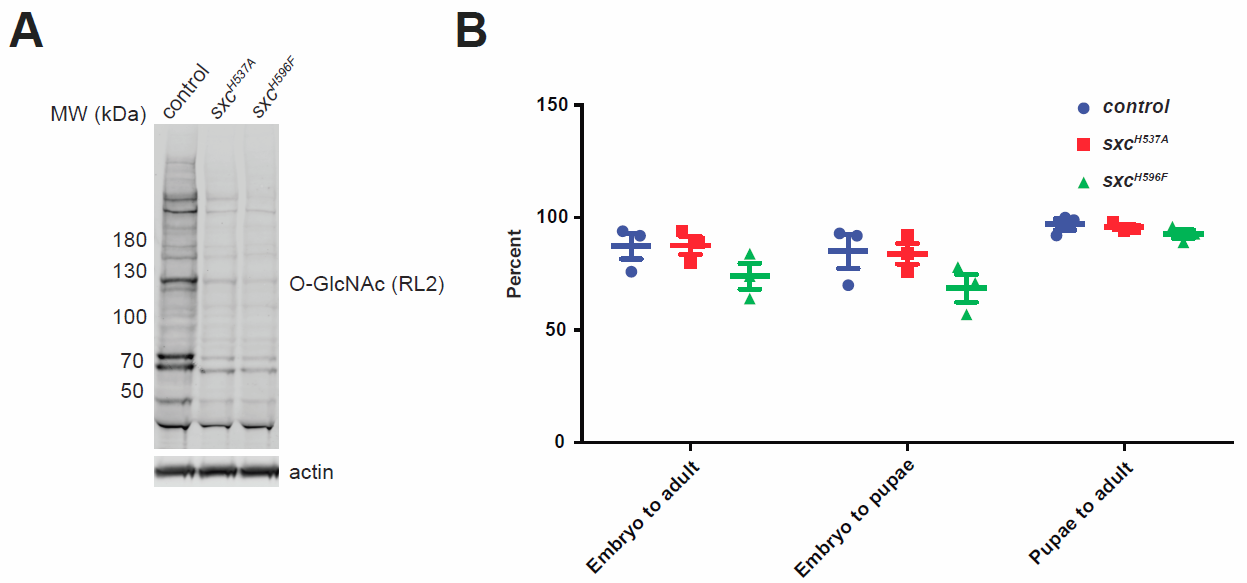

Fig. S4. Western Blot and developmental survival of *sxc^H596F^* flies

(A) Embryos from either wildtype, *sxc^H537A^*, *sxc^H596F^* homozygotes were assessed for levels of global O-GlcNAc using a pan-O-GlcNAc antibody RL2. The blot was normalized to actin. This blot is a representative of three experiments.
(B) Reduced total O-GlcNAc levels in *sxc^H596F^* and *sxc^H537A^* homozygotes are not associated with developmental lethality. Data presented as percentage of pupae and adults derived from stage 11-16 embryos (100 per genotype per experiment, n=3). Based on Student’s t-test with Holm-Sidak’s correction for multiple testing. Complete list of p-values and summary statistics is provided in Table S2.

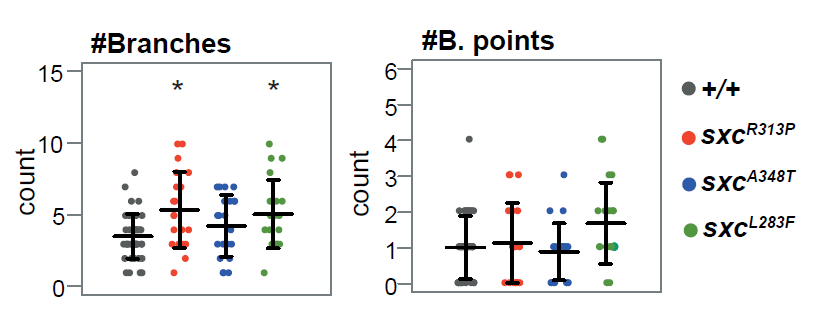

Fig. S5. NMJ branching morphology of *sxc^R313P^*, *sxc^A348T^* and *sxc^L283F^*

Number of synaptic branches is increased in *sxc^R313P^* (N = 21, *p* = 0.049, in red) and *sxc^L283F^* larvae (N = 20, in green) compared to the genetic background control (*+/+*, N = 41, in grey). Number of branching points is not significantly different. Data presented as individual data points with mean ± SD. * p< 0.05. P-values are based on Kruskal-Wallis test with Wilcoxon pairwise test for multiple comparisons. Complete list of p-values and summary statistics is provided in Table S3.

Table S1. Primer sequences.

| **Line** | **Purpose** | **Primer name** | **Sequence** |
| --- | --- | --- | --- |
| H596F | Guide RNA | CF_DmT596_fwd | GTCG TTATCTTAGCTCGGACTTC |
| H596F | Guide RNA | CF_DmT596_rev | AAAC GAAGTCCGAGCTAAGATAA |
| H596F | Mutagenesis | H596F wobble F | CTTCGAATAGGTTATCTTAGCTCGGAtTTtGGcAATttTCCTACCTCGCATTTAATGCAATCTG |
| H596F | Mutagenesis | H596F wobble R | CAGATTGCATTAAATGCGAGGTAGGAaaATTgCCaAAaTCCGAGCTAAGATAACCTATTCGAAG |
| H596F | Diagnosis | T59 DIG F | ATTCAGTCTTACCGAACGGCTCTAAAG |
| H596F | Diagnosis | T59 DIG R | GACTCTCGACTGATTTTGTGTCGAAATG |
| H596F | Sequencing | T59 DIG SEQ | CCGTTTACCATCCGTGCATC |
| L283F | Guide RNA | CF_DmT283_F | GTCG CCATGAACTACAGCATTATT |
| L283F | Guide RNA | CF_DmT283_R | AAAC AATAATGCTGTAGTTCATGG |
| L283F | 2kb PCR | T2fix_F | aaaGGATCc acacgaaagttttgggagccctttgg |
| L283F | 2kb PCR | T2fix_R | tttGCGGCCGC ATTCCCGAAGTCCGAGCTAAGATAACC |
| L283F | RF mutagenesis | L283F wobble R | GCCGCCTATTTACGTGCATTAAATtTcTCTCCGAAcAAcGCcGTgGTTCATGGCAATTTGGCATGCGTTTAC |
| L283F | RF mutagenesis | L283F wobble R friend | GCCCCGCTGACATCTTGCAATTC |
| L283F | Diagnosis | T2 DIG F | ccagttaaaggttttatggaaaaggtacg |
| L283F | Diagnosis | T2 DIG R | GCGTAGCTTCTTCAATATACCCCTGC |
| L283F  R313P  A348T | Line check | T2 OOB F  (283, 313 + 348) | Cggtaccccttacaaagatagtgagag |
| L283F | Line check | T2 OOB R (283) | CACAGATTGCATTAAATGCGAGGTAGG |
| R313P | Guide RNA | CF_DmT313_fwd | GTCG CAGTACGCATCGGGGAAATT |
| R313P | Guide RNA | CF_DmT313_rev | AAAC AATTTCCCCGATGCGTACTG |
| R313P | 2kb PCR | R313 FIX F | AAAggatcc Gatagaatgaaaaaatgaagtctttagaatc |
| R313P | 2kb PCR | R313 FIX R | TTTGCGGCCGC TACCATCATCTGGACTTAAGGCATAGC |
| R313P | RF mutagenesis | R313 wobble F (and T2 DIG R pair) | CTTATTGATTTAGCTATCGATACATATAGG**ccc**GCAATAGAACTGCAACCAAA**c**TT**t**CC**g**GA**c**GCGTACTGCAATCTTGCAAACGC |
| R313P | Diagnosis | DmT R313 DIG F | agttgaaatgtacaagcaacgtcagaac |
| R313P | Diagnosis | DmT R313 DIG R | CAGCTTCTTTAACctataaagattacttag |
| A348T | Guide RNA | CF_DmT348_fwd | GTCG tttatagGTTAAAGAAGCTG |
| A348T | Guide RNA | CF_DmT348_rev | AAAC CAGCTTCTTTAACctataaa |
| A348T | 2kb PCR | T348 BA F | aaaGGATCC Tattgtagttgggaaattggactataacg |
| A348T | 2kb PCR | T348 BA R | aaaGCGGCCGC ACTTGAATAGGAGCAGGGCGAAGAGC |
| A348T | RF mutagenesis | T348 patcher F (with L283F wobble R friend) | gtgcactaactaagtaatctttatagGTgAAgGAgGCcGAGGACTGCTATAACACTaCcCTAAGACTATGTTCAAATCATGCAGATTC |
| A348T | Sequencing | T348 seq 1 | Ttttgaaactaacccttcacac |
| A348T | Sequencing | T348 seq 2 | AAAGATTCCGGAAATATTCCTG |
| A348T | Line check | T348 OOB R (common line primer with R313P) | CTAGCCCCACTAGTACCTGGATAGC |

Table S2. Light-off jump habituation parameters summary

| **Condition** | **N total** | **N jumpers** | **%**  **of jumpers** | **mean TTC** | **mean TTC SD** | **mean**  **TTC SEM** |
| --- | --- | --- | --- | --- | --- | --- |
| contro:l +/+ (Figure 1A) | 96 | 65 | 67% | 4,4 | 0,9 | 0,4 |
| Sxc^H537A/+^ (Figure 1A) | 96 | 59 | 61% | 12,8 | 6,7 | 2,7 |
| control: +/+ (Figure 1B) | 96 | 72 | 75% | 9,4 | 3,5 | 1,4 |
| Sxc^H537A/+^ (Figure 1B) | 96 | 72 | 75% | 15,5 | 7,8 | 3,2 |
| Oga^KO/+^ (Figure 1B) | 96 | 76 | 79% | 17,1 | 10,4 | 4,2 |
| Sxc^H537A/+^; Oga^KO/+^ (Figure 1B) | 96 | 70 | 73% | 10,2 | 5,9 | 2,4 |
| control: elav-Gal4/+  (Figure 1C) | 64 | 38 | 59% | 2,6 | 1,3 | 0,7 |
| elav-Gal4>UAS-sxc  (Figure 1C) | 64 | 55 | 85% | 12,8 | 4 | 2 |
| control: +/+ (Figure 1D) | 96 | 65 | 68% | 4,4 | 0,9 | 0,4 |
| sxc^H537A/+^; elav-Gal4/+  (Figure 1D) | 64 | 40 | 63% | 14,6 | 6 | 3 |
| sxc^H537A/+^; elav-Gal4>UAS-sxc (Figure 1D) | 64 | 43 | 67% | 4,2 | 1,3 | 0,7 |
| control: +/+ (Figure 4A,B) | 96 | 65 | 68% | 4,4 | 0,9 | 0,4 |
| sxc^R313P/+^ (Figure 4A) | 96 | 73 | 76% | 12,7 | 14,9 | 2 |
| sxc^A348T/+^ (Figure 4B) | 96 | 72 | 75% | 8,6 | 4,5 | 1,8 |
| sxc^A348T/A348T^ (Figure 4B) | 96 | 76 | 80% | 24,1 | 3,9 | 1,6 |
| control: (Figure 4C,E) | 96 | 74 | 77% | 7,4 | 2,4 | 1 |
| sxc^R313P/+^ (Figure 4C,E) | 96 | 81 | 84% | 27,3 | 12,2 | 5 |
| Oga^KO/+^ (Figure 4C) | 96 | 58 | 60% | 7,3 | 3,4 | 1,4 |
| sxc^R313P/+^; Oga^KO/+^ (Figure 4C) | 96 | 53 | 56% | 10,6 | 6,5 | 2,6 |
| Oga^D133N/+^ (Figure 4E) | 96 | 69 | 72% | 14,9 | 11,8 | 4,9 |
| sxc^R313P/+^; Oga^D133N/+^  (Figure 4E) | 96 | 64 | 67% | 9 | 5,4 | 2,2 |
| control: +/+ (Figure 4D,F) | 96 | 72 | 75% | 9,4 | 3,5 | 1,4 |
| sxc^A348T/A348T^ (Figure 4D,F) | 96 | 79 | 82% | 33,5 | 10 | 4,1 |
| Oga^KO/KO^ (Figure 4D) | 96 | 87 | 90% | 25,6 | 9,3 | 3,8 |
| sxc^A348T/A348T^; Oga^KO/KO^  (Figure 4D) | 96 | 62 | 65% | 14,2 | 5 | 2 |
| Oga^D133N/D133N^ (Figure 4F) | 96 | 89 | 93% | 57,8 | 18,7 | 7,6 |
| sxc^A348T/A348T^; Oga^D133N/D133N^ (Figure 4F) | 96 | 56 | 58% | 8,2 | 3,5 | 1,4 |
| Sxc^H537A/H537A^ (not depicted) | 64 | 23 | 36% | NA | NA | NA |
| elav-Gal4>UAS-sxc^RNAi^  (not depicted) | 96 | 18 | 19% | NA | NA | NA |
| sxc^R313P/R313P^ (not depicted) | 64 | 31 | 48% | NA | NA | NA |
| sxc^L283F/+^ (not depicted) | 96 | 44 | 46% | NA | NA | NA |
| sxc^L283F/L283F^ (not depicted) | 96 | 38 | 40% | NA | NA | NA |

Table S3 (separate file). Summary statistics

Table S4 (separate file). Enrichment in *Drosophila* phenotypes and orthologs of human genes implicated in intellectual disability
